# A new automated chilled adult release system for the aerial distribution of sterile male tsetse flies

**DOI:** 10.1101/2020.04.14.040915

**Authors:** Caroline K. Mirieri, Gratian N. Mutika, Jimmy Bruno, Momar Talla Seck, Baba Sall, Andrew G. Parker, Monique M. van Oers, Marc J.B. Vreysen, Jeremy Bouyer, Adly M. M. Abd-Alla

## Abstract

**Background:** Tsetse flies transmit trypanosomes that cause the debilitating diseases human African trypanosomosis (HAT) or sleeping sickness in humans and animal African trypanosomosis (AAT) or nagana in livestock. The riverine tsetse species *Glossina palpalis gambiensis* Vanderplank (Diptera: Glossinidae) inhabits riparian forests along river systems in West Africa. The Government of Senegal has embarked on a project to eliminate a population of this tsetse species from the Niayes area with the objective to manage AAT in this area. The project is implemented following an area-wide integrated pest management approach with an SIT component. The SIT can only be successful when the sterile males that are released in the field are of high biological quality, i.e. have the same dispersal capacity, survival and competitiveness as their wild counterparts. To date, sterile tsetse males have been released by air using biodegradable cardboard cartons that were manually dropped from a fixed-wing aircraft or gyrocopter. The cardboard boxes are however expensive, and the system is rather cumbersome to implement.

**Methods:** A new prototype of an automated chilled adult release system (Bruno Spreader Innovation, (BSI™)) for tsetse flies was tested for its accuracy (in counting numbers of sterile males as loaded into the machine), release rate consistency and impact on quality of the released males. The impact of the release process was evaluated on several performance indicators of the irradiated male flies such as flight propensity, survival, mating competitiveness, premating and mating duration, and insemination rate of mated females.

**Results:** The BSI™ release system counted with a consistent accuracy and released homogenously tsetse flies at the lowest motor speed (0.6 rpm). In addition, the chilling conditions (6 ± 1 °C) and the release process (passing of flies through the machine) had no significant negative impact on the males’ flight propensity. No significant differences were observed between the control males (no irradiation and no exposure to the release process), irradiated males (no exposure to the release process) and irradiated males exposed to the release process with respect to mating competitiveness, premating period and mating duration. Only survival of irradiated males that were exposed to the release process was reduced (∼2 folds), irrespective of whether the males were held with or without feeding.

**Conclusion:** Although the release process had a negative effect on survival of the flies, the data of the experiments indicate that the BSI machine holds promise for use in tsetse SIT programmes. The results of this study will now need to be confirmed under operational field conditions in West Africa.

## Introduction

Tsetse flies (*Glossina* spp.; Diptera: Glossinidae) are hematophagous insects and are the cyclical vectors of two debilitating diseases in sub-Saharan Africa, i.e. human African trypanosomosis (HAT) or sleeping sickness in humans and animal African trypanosomosis (AAT) or nagana in livestock [1,2]. Nagana and sleeping sickness have been major obstacles to rural development and a severe constraint for the development of more efficient and sustainable agricultural production systems in sub-Saharan Africa [3]. AAT limits the exploitation of fertile agricultural land in ∼10 million km^2^ in this part of Africa and, therefore, tsetse flies are rightly considered one of the root causes of poverty and hunger [4,5]. In West Africa, tsetse flies of the *palpalis* group, i.e. *Glossina palpalis palpalis* (Robineau-Desvoidy), *Glossina palpalis gambiensis* Vanderplank and *Glossina tachinoides* Westwood, are the most important cyclical vectors of these two diseases [6].

Due to the lack of effective vaccines and inexpensive drugs for HAT, and the development of resistance of the AAT parasites against available trypanocidal drugs [7], tsetse control remains a key component for the integrated sustainable management of both diseases [6]. Currently, there are four acceptable control tactics for the integrated management of tsetse vectors, i.e. (i) the live-bait technique (dip, spray, or pour on application of residual insecticides on livestock) [8], (ii) insecticide-impregnated targets/traps [9], (iii) the sequential aerosol technique (SAT) [10], and (iv) the sterile insect technique (SIT) [11–13]. In most cases, sustainable management of tsetse fly populations can only be achieved if the control tactics are implemented following the principles of area-wide integrated pest management (AW-IPM) [14]. For example, IPM applied against an entire pest population within a delimited geographic area, with a minimum size large enough or protected by a buffer zone so that natural dispersal of the population occurs only within this area [15].

Already in the 1970’s, an attempt was made to eradicate the *G. p. gambiensis* population from the Niayes region in Senegal, mainly using insecticide-based control tactics. However, area-wide principles were not adhered to and the project failed to create a sustainable tsetse-free zone, leading to re-colonization of the Niayes from relict pockets that had not been treated [16–18].

In 2005, the Government of Senegal embarked on a campaign to eradicate a population of *G. p. gambiensis* [16] from a 1,000-km^2^ area of the Niayes, located in the vicinity of the capital Dakar. The programme has been implemented under the auspices of the Pan African Tsetse and Trypanosomosis Eradication Campaign (PATTEC), a political initiative started in 2001 that calls for intensified efforts to reduce the tsetse and trypanosomosis problem [19]. The Direction des Services Vétérinaires (DSV) of the Ministry of Livestock (Ministère de l’Elevage et des Productions animales) implemented the eradication campaign with support from the Institut Sénégalais de Recherches Agricoles (ISRA) of the Ministry of Agriculture (Ministère de l’Agriculture et de l’Equipement rural). The programme received financial and technical support from the Food and Agriculture Organization of the United Nations (FAO), the International Atomic Energy Agency (IAEA), the Centre de Coopération Internationale en Recherche Agronomique pour le Développement (CIRAD), and the USA State Department under the Peaceful Uses Initiative (PUI). In this AW-IPM programme, conventional suppression methods (insecticide-impregnated traps/targets/nets and insecticide pour-on on livestock, insecticide ground and aerial spray, bush clearing) were integrated with the release of sterile male flies [20].

The sterile insect technique (SIT) is a species-specific, safe, efficient, environment-friendly autocidal control tactic to manage populations of insect pests and disease vectors [21–24]. The SIT requires mass-rearing of the target insect, sterilization of the males using ionizing radiation and sequential area-wide releases of a large numbers of the sterile males into the target area. The sterile male flies compete with wild male flies for mating with the wild female population, interrupting their reproductive potential and ultimately resulting in population reduction or elimination [25,26]. The released sterile males, therefore, need to have adequate mobility to find and mate with virgin wild females, and this is vital to successfully implement the SIT component of AW-IPM programmes [25,27,28]. Therefore, routine quality control procedures are crucial to identify weaknesses in fly production and handling procedures that result in low quality of the sterile males, as this may lead to potential programme failure [14,29].

The aerial release of sterile male tsetse flies was pioneered in the programme that eradicated a population of *Glossina austeni* from Unguja Island of Zanzibar in the 1990’s [13]. The sterile males were packaged in bio-degradable cartons that were manually dropped from fixed-wing aircrafts [30]. A similar approach was used during the initial years in the Senegal project, but the boxes were dropped from gyrocopters [20]. In view of the high cost of the bio-degradable cartons and the lack of storage space in the gyrocopters (requiring frequent landings to reload), efforts were made to develop chilled-adult release systems similar to those developed for the release of sterile fruit flies [31,32]. However, the release systems for these pests have a very high throughput to obtain release densities of 2,500 to 200,000 fruit flies/km^2^/week [33], but for tsetse flies the challenge was to develop a machine that could disperse the sterile insects at very low release rates to obtain densities of 15 – 80 flies/km^2^/week [33]. The aerial release of sterile insects has many advantages as compared with ground releases i.e. it is fast, reaches areas that are inaccessible for ground release and provides a uniform distribution of sterile insects over the target areas. However, it is expensive and in some larger programmes, it represents around 40% of the annual operating budget of the sterile fly emergence and release centres [34]. Therefore, attempts to reduce the cost of the aerial release process are desirable either by reducing the frequency of flights through increasing the quality of the sterile males [30,31] or by using automated chilled-adult release systems that avoid the cost of bio-degradable carton boxes [30,31]. Currently, the majority of SIT programmes release chilled adult insects into the targeted areas [34,35] using small fixed-wing air craft, helicopters or gyrocopters [30,34,36,37]. The use of smaller airplanes or gyrocopters is one way to reduce the aerial release cost, e.g. the use of a gyrocopter in Senegal was the cheapest way at a cost of € 320 per flying hour during the period 2010-2020 [2]. Similarly, replacing the paper boxes and reducing the number of flights using an automated chilled-adult release system will not only reduce the implementation cost but also increase the efficiency of the program through improved sterile male fly distribution [30,33].

The use of automated devices to release chilled male tsetse flies was earlier (in 2012 for mexicana 1 and 2015 for Mexicana 2) attempted in Senegal using a Mubarqui Smart Release Machine (MSRM2), adapted from the one used to release fruit flies (MSRM1) [37]. Initially, the system gave promising results demonstrating its potential suitability for use in tsetse SIT programmes and provided acceptable distribution when the males were maintained at 9-12°C. However, its operational use revealed its limitation in terms of inadequate consistency of the release rates, which was probably caused by vibrations of the gyrocopter that interfered with the vibrator of the MSRM2 that was used as a fly ejection mechanism [37]. This resulted in an average recapture rate that was lower than the one obtained when sterile male *G. austeni* were released at ambient temperature (25-29°C) using paper cartons in the Zanzibar programme [30]. The disappointing recapture rates of the released male flies prompted the programme to discontinue the use of the MSRM2. A major drawback was that the impact of the release process on the sterile male tsetse quality, including their ability to fly and to survive after passing through the MSRM2, was not well tested before its operational use in Senegal [37]. In this study, a new automated chilled adult BSI prototype (Bruno Spreader Innovation (BSI™), the Aerial Works Company (AEWO), St-Jean le Vieux, France) was tested for the release of sterile male *G. p. gambiensis* flies under laboratory conditions. The release system contains a cylinder rotating against a brush as an ejection mechanism [38] that enables the release of a low number of sterile insects per unit area and time. First, the machine was calibrated, and the consistency of the release rate determined. Thereafter, the impact of the release process on sterile male performance was assessed in terms of flight propensity, mating competitiveness (in walk-in field cages), premating period and mating duration, insemination potential and survival.

## Material and Methods

### Tsetse Flies

All experiments were carried out with flies from a *G. p. gambiensis* colony that was established in 2009 at the FAO/IAEA Insect Pest Control Laboratory (IPCL), Seibersdorf, Austria [39,40] from pupae received from a colony maintained at the Centre International de Recherche-Développement sur l’Elevage en zone Subhumide (CIRDES), Bobo Dioulasso, Burkina Faso. The original colony was established in 1972 at Maison-Alfort, France from field pupae collected in Guinguette, Burkina Faso and then transferred to CIRDES in 1975.

The colony at the IPCL has been maintained using an *in vitro* feeding system with thawed bovine blood (Svaman spol, s.r.o., Myjava, Slovak Republic). The blood was kept frozen at −20°C and irradiated with 1 kGy in a commercial 220 PBq ^60^Co wet storage panoramic shuffle irradiator. The flies were offered a blood meal three times a week and maintained under a 12L:12D light regime cycle. Pupae were incubated at 24.1 ± 0.1°C and 78.8 ± 3.7% R.H. for four weeks and adults emerged under the same temperature and humidity conditions. These conditions will henceforth be referred to as standard laboratory rearing conditions.

### Radiation

The tsetse puparia were irradiated in air at the IPCL, Seibersdorf, Austria using a Gammacell^®^ 220 (MDS Nordion Ltd., Ottawa, Canada) ^60^Co irradiator. The dose rate was measured by alanine dosimetry as 2.144 Gy·sec^− 1^ on 2015-03-03 with an expanded uncertainty (k = 2) of 3.2%. The radiation field was mapped using Gafchromic HD-V2 film and the dose uniformity ratio in the volume used for the experiments was < 1.1. The irradiated group was placed in a petri dish at the centre of a polycarbonate jar (2200 mL) and sandwiched between two phase change packs that kept the temperature below 10°C during irradiation with 120 Gy. Untreated puparia or flies were used as control (0 Gy) and handled in the same manner.

### BSI™ automated release device

The BSI™ automated chilled adult release system (hereafter referred to as “BSI™”, **Figure 1, Supplementary file 1**) and the associated software (BSI Navigator version 1.9.8) installed on a tablet computer (Samsung Galaxy Tab S2), were tested (flight simulations) at the IPCL for its functionality (accuracy and release rate consistency) and impact (chilling and potential physical damage) on the sterile adult males. The BSI™ has a weight of 20 kg and consists of a funnel surrounded by a cooling unit, into which the flies are loaded and held until discharged into cavities on a rotating cylinder that acts as an ejection mechanism, resulting in the release of flies [38]. An optical sensor monitors the number of males released. The machine was switched on at least one hour before the loading of the flies to obtain a stable temperature of 6 ± 1°C.

**Figure 1.**
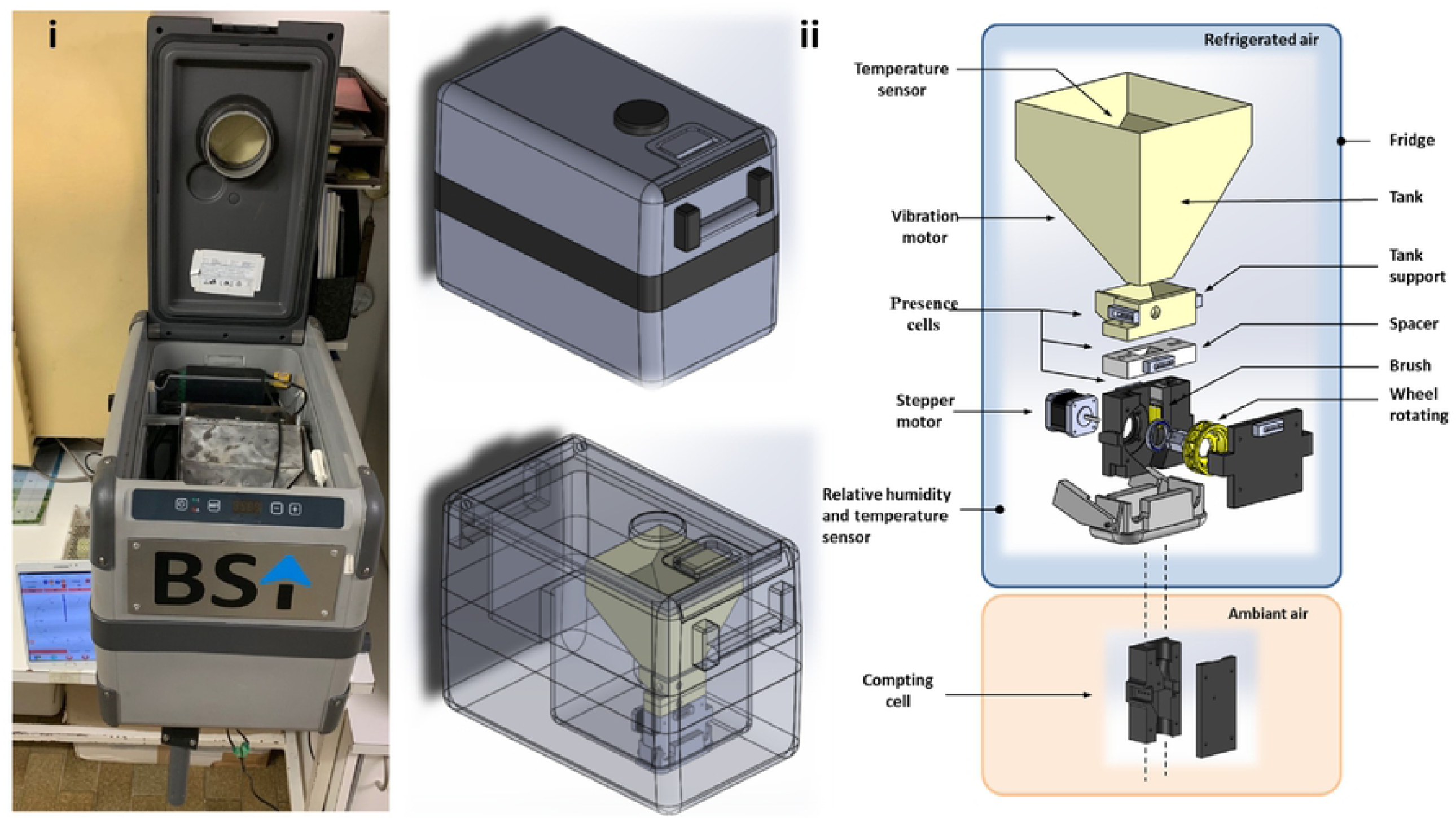
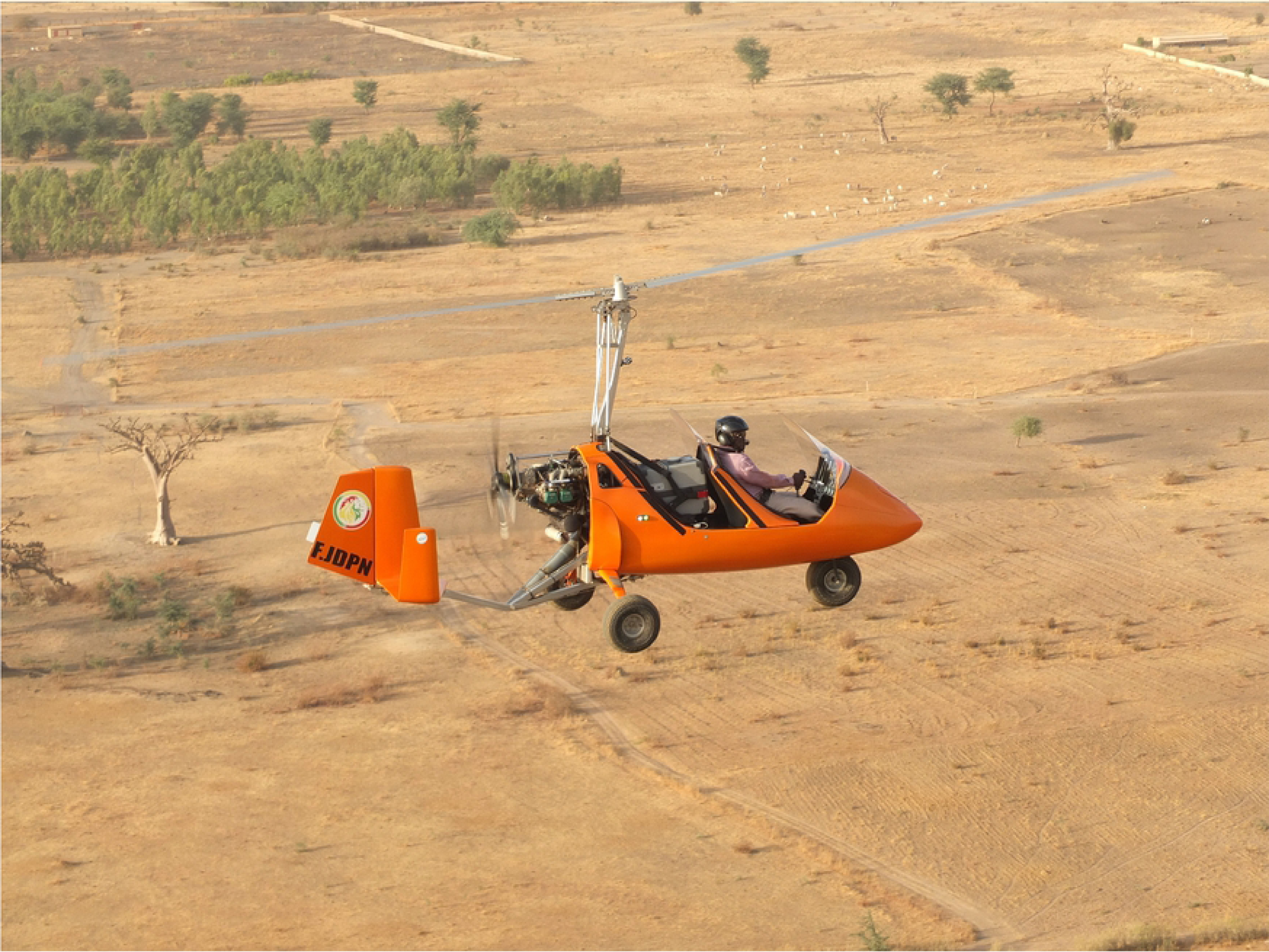
The BSI™ a: images showing (i) top view of the open BSI™ showing the storage funnel where tsetse flies are held at 6°C, (ii) 3-dimention technical drawing showing different parts on the BSI release machine. b: a gyrocopter carrying the BSI release machine in field operation.

### Calibration of the BSI™

The accuracy of the BSI™, i.e. its ability to count the numbers of flies released was assessed at speeds ranging from 0.6 to 6 rpm. After selecting a specific speed, each of three batches of a known number (480, 1000 or 2000) of 5-day old sterile male *G. p gambiensis* were immobilized by exposure to a gentle flow of cold air (4°C) in a chiller and loaded into the BSI™ and released. As the chiller and the BSI™ were located in different rooms, the chilled flies were transferred from the chiller to the BSI™ on a netted circular container placed on a plastic box filled with ice and covered with aluminium foil to keep the flies immobilized. The released flies were collected on ice to keep them in an anaesthetized state for repeated use in calibration at each rotation speed. Each of the three batches was replicated three times for each of the ten rotation speeds. The flies were discarded after the calibration. The BSI™ software counted and automatically recorded the number of sterile male flies passing through the machine as detected by an optical sensor. This figure was later compared with the hand counts of the flies loaded into the machine to calculate the error. The software automatically generated graphs with trend lines of error rate for the ten selected rotation speeds used in each replicate within the 3 batches of flies. This enabled the selection of the replicate with the best linear curve (with the least errors at each speed), which was later used for the simulated flight releases.

### Consistency of the release rate

During calibration the best accuracy was observed at a rotation speed of 0.6 rpm and was therefore used, to assess the consistency of the release rate of the BSI™, i.e. the number of males released per minute. A total of 2000 male flies were exposed to the release process at a rotation speed of 0.6 rpm for three replicates. The BSI™ was switched on one hour before the test and the software set to the following parameters for release of the flies: i) IPCL polygon where the machine was physically located, ii) manual control of the release iii) the numbers of insects available to pass through the machine, iv) a rotation speed of 0.6 rpm and v) a vibration value of the maximum power of the shaker. The duration of the release process was recorded using a digital timer. Male flies that passed through the BSI™ were collected in a container placed on ice. The collection containers were changed every five minutes and the males counted manually. These manual counts were used for comparison with the number of males recorded by the BSI™. A total of seven containers of released flies were collected for each replicate. The flies were discarded after the test.

### Impact of the release process on sterile male performance

#### Preparation of the tsetse flies

After emergence of the female *G. p. gambiensis* flies, the remaining male pupae were collected and divided into two groups. Both groups were chilled at 10°C for one hour before one group of pupae was irradiated. Thereafter, both groups of pupae were covered with sieved, sterilized sand mixed with 0.025% fluorescent powder by mass following the procedure used in the operational program in Senegal [41]. Different colours were used for the irradiated and untreated groups. The pupae were placed in emergence cages and the emerged marked males were maintained under standard colony conditions. Virgin female flies were collected from the colony three days prior to the mating competitiveness test. For this study, five different treatment groups of males were used: irradiated males exposed to the release process that were held in the machine for 5 minutes (5m), 60 minutes (60m) and 120 minutes (120m) before passing through the machine, irradiated males that were not exposed to the release process (Irrd) and males from the colony that were neither irradiated nor exposed to the release process (Control). All male flies were offered two blood meals on the 1^st^ and 3^rd^ day after emergence, and 24 hours after the last blood meal, 5-day old males of the treatment groups were immobilised at 4°C for 18 and 36 minutes to allow sorting and counting. A total of 500 male flies were placed into the BSI™ to assess their flight propensity and mating competitiveness and survival after passing them through the machine after 5, 60 and 120 minutes compared with irradiated only and control.

#### Flight Propensity

The flight propensity of the 5-day old males of the treatment groups was assessed under standard rearing conditions. Flight tests were carried out following the modified FAO/IAEA/USDA protocol [42] in netted cages (45 × 45 × 45 cm) containing a black painted PMMA flight tube (89 mm diameter, 3 mm thick wall, 100 mm high). Light could only enter the tube from the top and the walls were coated with unscented talcum powder to prevent the flies from crawling out of the tube [40]. For each test, an average of 140 (49-410) chilled males were put in a plastic Petri dish (90 mm diameter) with the base covered by black porous paper and the flight tube placed on top. The number of male flies that escaped from the tube (called “flyers”) and those that remained in the tubes (called “non-flyers”) were recorded after two hours [43]. The cages were placed in a room with fluorescent lights giving an intensity of 500 lux at the flight tubes to attract the emerged flies. Each treatment was replicated eight times. Samples of 30 and 120 male flies that escaped the tube were collected for each of the treatment groups and the control group, respectively, for use in the mating competitiveness test seven days post emergence. The remainder of the flies were discarded.

#### Mating competitiveness and insemination rate

Mating competitiveness of the male flies was assessed in walk-in field cages containing a potted tree to simulate a natural environment. The netted cylindrical field cages [44] (2.9 m diameter and 2.0 m high) [45] were located inside a greenhouse with temperature and humidity control and natural light that could be supplemented with artificial light from cold white fluorescent tubes. The temperature in the greenhouse ranged from 24°C to 31°C and the relative humidity from 41 to 56% during the observation periods. Light intensity varied from 236 to 5000 lux depending on the position in the cage with areas under the PVC supporting frame and tree leaves recording lower light intensity. Temperature and humidity were recorded from 08:45 h am to 11:30 h am.

All mating competitiveness tests were carried out between 9.00 h am and 11.00 h am as described in previous experiments [46]. The male flies of the five treatment groups that were flyers from the flight propensity test, were released in the field cage to compete for mating opportunities with untreated colony males for mating with virgin females. The competitiveness test of each treatment group was replicated eight times. During the test in each replication, four field cages representing the four treatments were located in the centre of the greenhouse and used simultaneously for observations of mating activity, with each of two observers managing two cages.

In each field cage 30 three-day old virgin females were released ten minutes before two groups of 7-dayold flies (30 males from the control and 30 males from one of the four treatments, as previously described) to compete for mating opportunities at an initial ratio of 2 males:1 female, during the 2-hour observation. The male fly treatments were randomly allocated to different cages each day such that at the end of the experiment each treatment was observed in the same cage at least twice **(Table 2)**. When the male had successfully engaged the female *in copula*, the mating pair was collected in a tube with netting at both ends (4 cm diameter x 6 cm height) and after the couples disengaged, the males were separated from the females. Each tube was numbered to identify the individual male and female treatment. The period between the release of the male flies in the field cage until copulation was recorded as the pre-mating time. The difference in time between the initiation of successful copulation and separation was recorded as the mating duration. After the end of the mating, males and females were separated to identify the male treatment and the females dissected to estimate spermathecal fill. The fluorescent dyes to discriminate male fly treatments were differentiated using a USB digital microscope (AM4113FVT2, Dino-Lite Europe, Almere, The Netherlands) with UV-light, connected to a PC for display. The female flies were dissected in physiological saline solution under a binocular microscope and the insemination rate and spermathecal fill were assessed subjectively at x100 magnification using a Carl Zeiss compound microscope connected to a PC for display [23]. The spermathecal fill was scored to the nearest quarter for each spermatheca separately as empty (0), partially-full (0.25, 0.50 or 0.75) or full (1.0) and the quantity of sperm transferred was then computed as the sum of the two spermathecal scores. The number of females that mated as a proportion of the total females in each replicate is an indication of the tendency of the flies to mate (proportion mating, PM). The relative mating index (RMI) was defined as the number of mating pairs accounted for by the treatment category as a proportion of the total number of mating pairs. These indices represent the competitiveness of treated males relative to the colony control males [46].

**Table 1:** Averages (±SE) counts of calibration of the BSi release machine using batches of 480, 1000 and 2000 *G. palpalis gambiensis* males

| Speed (rpm) | 0.6 | 1.2 | 1.8 | 2.4 | 3.0 | 3.6 | 4.2 | 4.8 | 5.4 | 6.0 |
| --- | --- | --- | --- | --- | --- | --- | --- | --- | --- | --- |
| **counts ±SE (480)** | 252.7 ± 7.2 | 224.7 ± 4.4 | 213.7 ± 4.3 | 218.7 ± 6.1 | 199 ± 8.5 | 190 ± 3.1 | 186.7 ± 5.8 | 190.7 ± 5.8 | 175.7 ± 8.8 | 159.7 ± 2.9 |
| % error | 47.4 ± 1.5 | 53.2 ± 0.9 | 55.5 ± 0.9 | 54.4 ± 1.2 | 58.5 ± 1.9 | 60.4 ± 0.8 | 61.1 ± 1.2 | 60.3 ± 1.3 | 63.4 ± 1.9 | 66.7 ± 0.8 |
| **counts ±SE (1000)** | 520.7 ± 6.6 | 476.3 ± 4.3 | 474.3 ± 19.6 | 428.3 ± 6.4 | 398.7 ± 10.8 | 380.3 ± 12.7 | 353.7 ± 13.0 | 348.3 ± 9.0 | 337.7 ± 9.1 | 319 ± 7.0 |
| % error | 47.9 ± 0.7 | 52.4 ± 0.2 | 52.6 ± 2.2 | 57.2 ± 0.7 | 60.1 ± 1.1 | 62 ± 1.2 | 64.6 ± 1.1 | 65.2 ± 0.6 | 66.2 ± 0.2 | 68.1 ± 0.7 |
| **counts ±SE (2000)** | 1082.7 ± 7.2 | 1009 ± 28.3 | 941 ± 30.0 | 871.7 ± 22.7 | 847.3 ± 3.2 | 823 ± 38.8 | 749.3 ± 11.9 | 755.7 ± 5.8 | 752.7 ± 16.9 | 695.7 ± 8.7 |
| % error | 45.9 ± 0.4 | 49.6 ± 1.4 | 53 ± 1.5 | 56.4 ± 1.1 | 57.6 ± 0.2 | 58.9 ± 1.9 | 62.5 ± 0.6 | 62.2 ± 0.3 | 62.4 ± 0.8 | 65.2 ± 0.4 |

\*Each combination of density and speed was technically replicated three times.

**Table 2.** The mean (±sd) response variables for the flight propensity test and mating competitiveness observations. 5m, 60m, and 120m: irradiated males held chilled for 5 min, 60 min and 120 min, respectively before passing through the BSI release machine, Irrd: irradiated males but not passed through the BSI release machine and Control: non-irradiated males from the colony.

|  | 5m | 60m | 120m | Irrd | Control |
| --- | --- | --- | --- | --- | --- |
| Flight propensity (%) | 77.8±8.61 | 75.7±7.2 | 73.9±12.1 | 77.2±6.3 | 88.3±7.3* |
| Relative mating Index | 0.46±0.14 | 0.46±0.11 | 0.41±0.10 | 0.39±0.16 | 0.56. ±0.14 |
| Premating time (minutes) | 45.00±39.00 | 33.00±30.00 | 37.00±35.00 | 45.00±39.00 | 40.00±34.00 |
| Mating duration (minutes) | 81.00±31.00 | 71.00±30.00 | 72.00±27.00 | 70.00±28.00 | 80.00±30.00* |
| Spermathecal Fill | 1.53(±0.78) | 1.36(±0.80) | 1.35(±0.84) | 1.53(±0.79) | 1.54(±0.73) |

\*: Treatment with significant difference from other treatments

#### Survival

To determine the impact of the release process on the longevity of irradiated males, a survival test was carried out with one group of flies kept under starvation and a second group kept under standard blood feeding conditions. Five-day old males (n = 30) from the treatment and control groups were placed in adult holding cages (diameter 110 mm x height 50 mm) and maintained under standard tsetse rearing conditions. Two groups of flies originating from two different batches of flies (from different weeks) were used for the treatments, resulting in two biological/true replicates. Within each group, treatments were also divided into two replicates resulting in two technical replicates. This totalled 4 replicates (2 biological and 2 technical replicates). Male mortality was recorded daily under starvation conditions and three times per week for the standard feeding conditions.

### Data analysis

The consistency of release rate data was analyzed using a linear mixed-effects model fitted by maximum likelihood. The flight propensity, the relative mating index and the spermatheca values data were analyzed using a generalized linear mixed-effects model fitted by maximum likelihood (Laplace approximation) with a “binomial (logit)” family. Treatments were used as fixed effects and replicates were used as random effects. The premating and mating duration data were analyzed using linear regression models. The number of matings achieved was tested for equality of performance between treated and control males using the RMI and the Log likelyhood ratio test for comparison of means [47]. Spermathecal fill categories were analyzed using a Kruskall-Wallis test [47,48]. The survival of flies of different treatments was analyzed using Kaplan-Meier survival curves [49]. Survival curves were compared using the cox.surv model where the treatment was used as explanatory variable and the survival as the response variable. The data were statistically analyzed and graphs created in Excel and R version 3.6.2. [50] using RStudio Desktop version 1.2.5033 [51] with the packages; ggplot2 [52], nlme [53], lme4 [54], survival [55], coxme [56] and MASS [57].

## Results

### Calibration of the BSI™

The calibration results were used to assess the accuracy of the machine by counting the loaded number of flies for comparison with the number of flies released as counted and recorded by the machine. The software automatically generated tables and graphs of the machine counts and calculated the error rates for the three batches (480, 1000, 2000 flies) and their replicates. The results showed average error rates above 45% of the numbers counted by the machine compared with the hand-loaded number of flies into the machine, and this error rate increased with increasing motor speed (F = 494, df = 1, 88, *P* < 0.001) (**Figure 2A**). The lowest error rates (47.4% ± 1.5, 47.9 ± 0.7; and 45.9 ± 0.4 for loads of 480, 1000 and 2000 flies, respectively) were observed at the lowest motor speed of 0.6 rpm. The error rate was not affected by the number of flies loaded into the machine (*P* = 0.245) (**Table 1**; **Figure 2B**).

**Figure 2:**
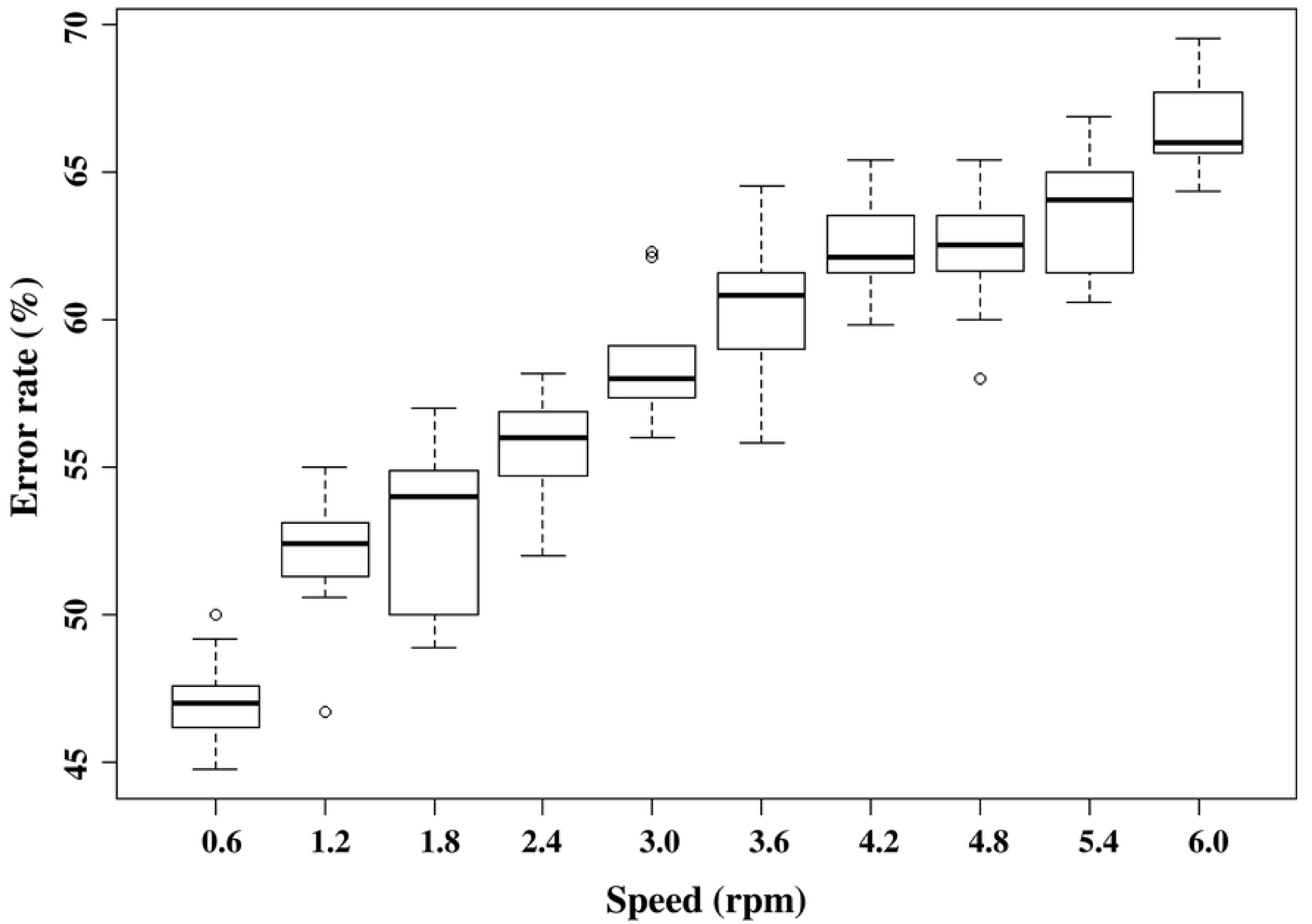
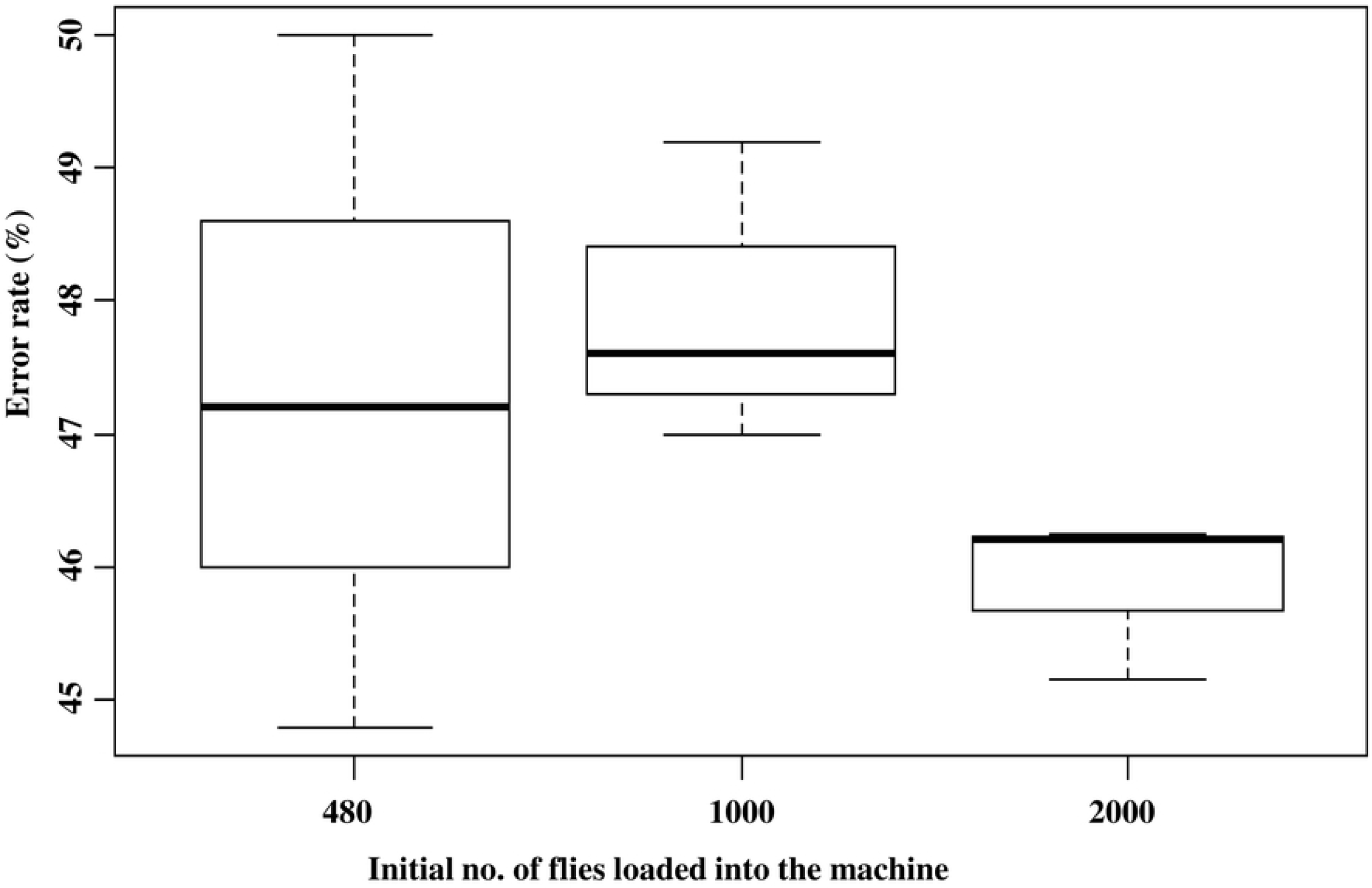
Effect of motor speed and initial fly number on the error rate of the BSI™. a: Impact of motor speed (rpm) on the error rate during counting of the sterile males released by the BSI™. b: Impact of initial number of flies on the error rate during the counting of sterile males released by the BSI™ at the lowest speed (0.6 rpm). The graphs represent the minimum, first quartile, median, third quartile and maximum for each treatment. Values indicated by the same lower-case letter do not differ significantly at the 5% level.

### Consistency of the release rate of the BSI™

The release rates of the sterile flies over time as shown by the manual hand-counts, the counts recorded by the optical sensor and estimated counts (after correction by the software using the error rate estimated as explained above) are presented in **Figure 3A**. The actual release rate as given by the manual counts did not vary significantly with time (F = 3.5849, df = 1, 20, *P* = 0.0736) after an approximately one-minute initial delay **(Figure 3B, Supplementary file 2**). There were no significant differences in release rates between the three replicates of the 2000 flies used (*P* > 0.05). The manual and estimated counts were strongly correlated (t = 67.348, df = 1, 22, *P* > 0.001) showing that the machine was able to correct any errors in counts during the release period, in an accurate and consistent manner (**Figure 3C**).

**Figure 3:**
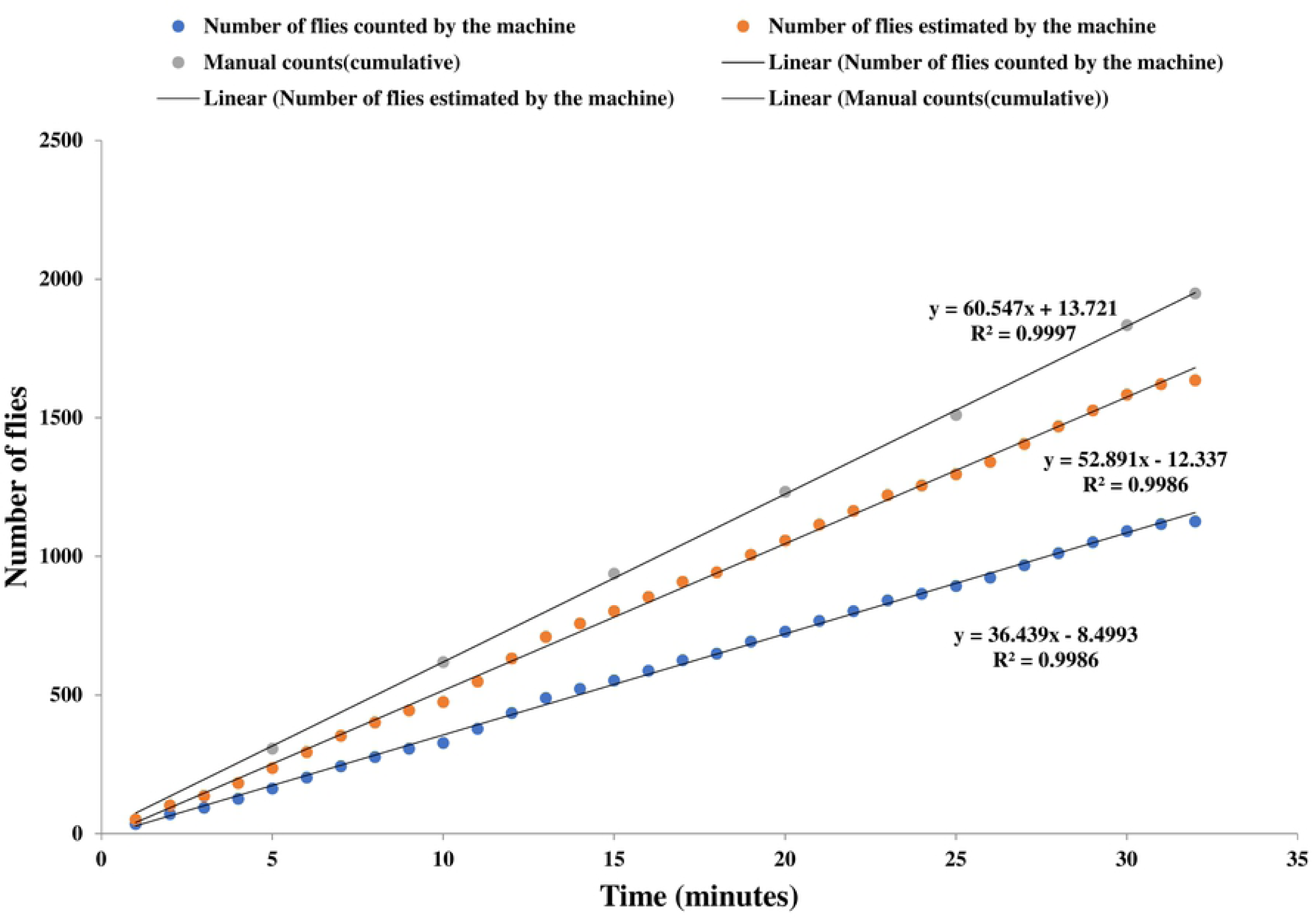
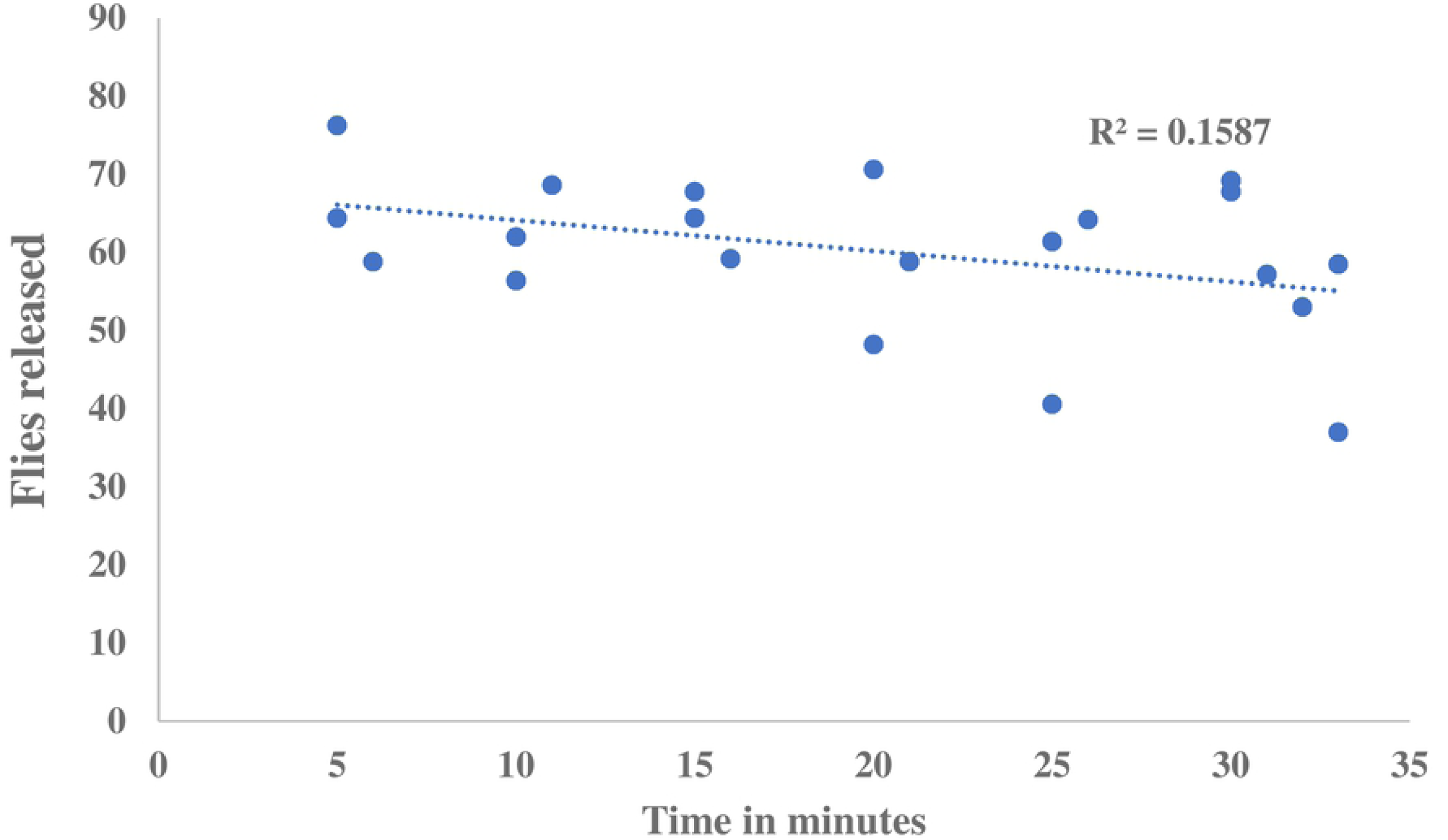
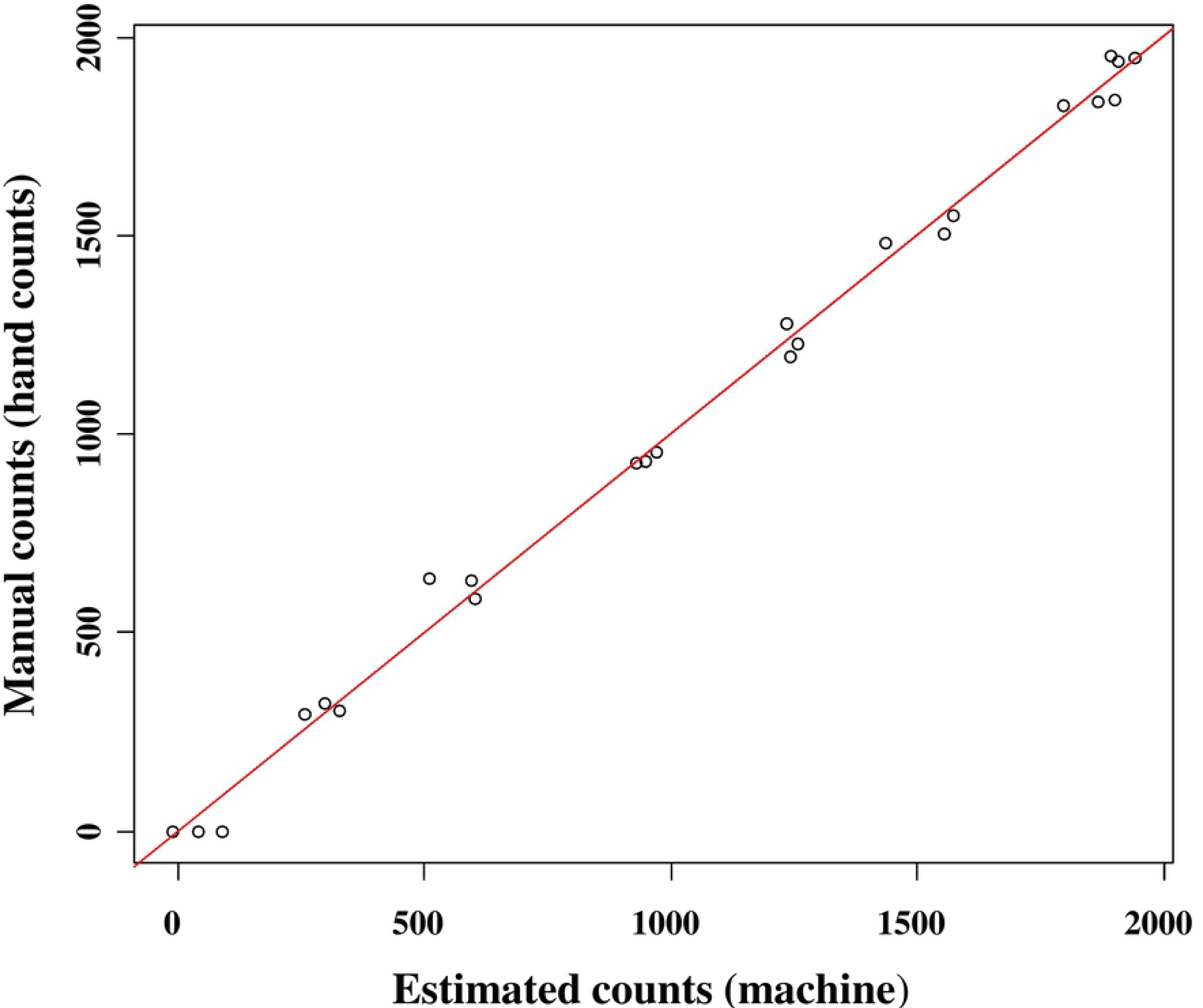
Consistency of release rate of BSI™ a: Cumulative count of flies (manual, actual counts and estimated counts by the machine) over time at the lowest speed (0.6 rpm). Comparison of the recorded release rate against the actual rate (by hand count) b: The release consistency of flies per minute, c; Prediction of manual counts from estimated counts using a linear model.

### Sterile male *G. p. gambiensis* performance after exposure to the release process with the BSI

#### a) Male flight propensity

The flight propensity of the male flies was more than 60% for all replicates and mating activity was observed in all cages for all treatments (**Table 2**). Untreated colony males had an average flight propensity of 88.3%, which was significantly higher than that of the treatment males (z = 5.290, *P* < 0.001). However, the flight propensity of males that were only irradiated (Irrd) and of irradiated males that were exposed to the release process for different durations (5, 60 and 120 minutes) was similar (**Figure 4, Table 2**).

**Figure 4:**
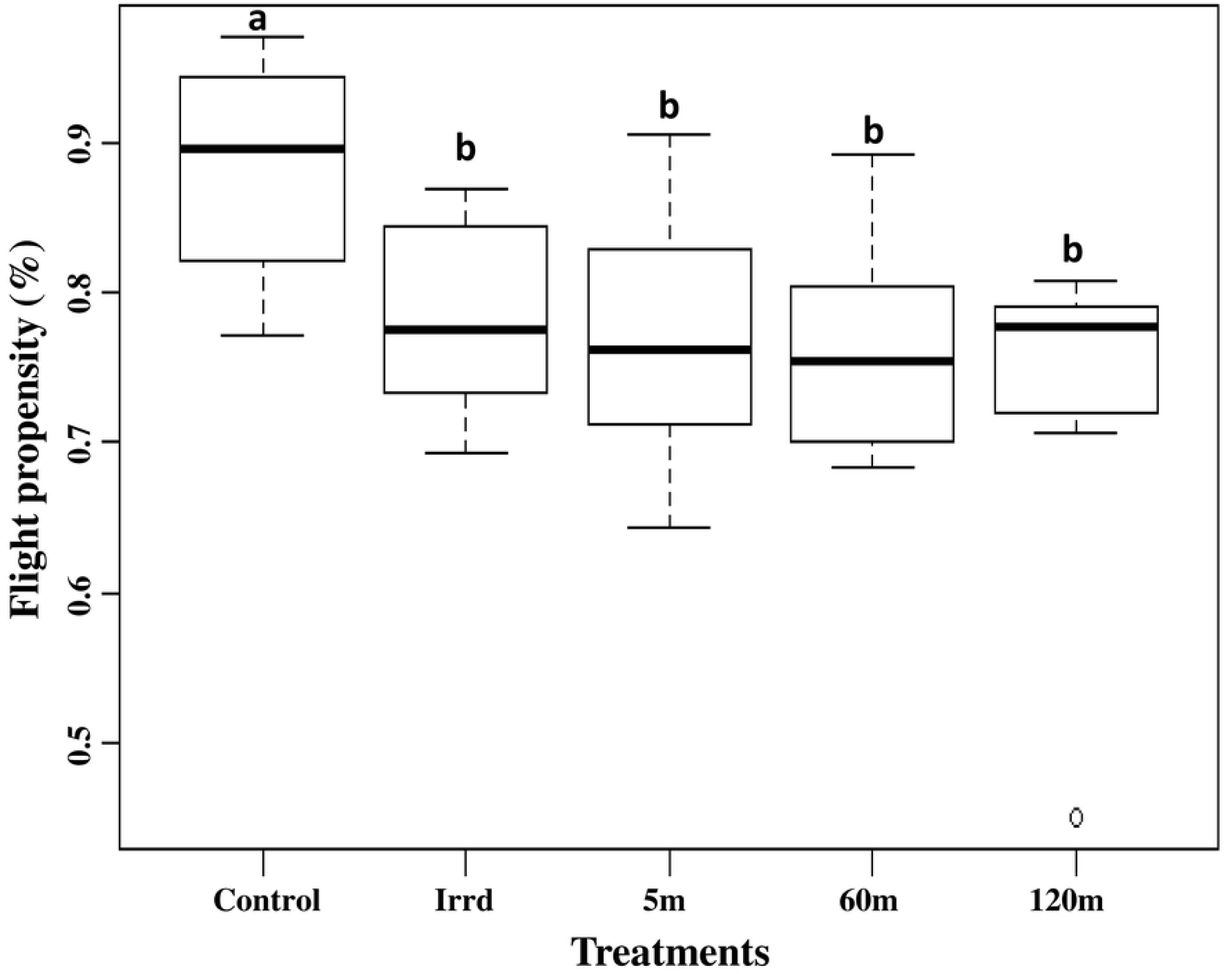
Impact of the release process through the BSI™ on male flight propensity. The graph represents the minimum, first quartile, median, third quartile and maximum for each treatment. Values indicated by the same lower-case letter do not differ significantly at the 5% level.

#### b) Mating competitiveness

When flies were released in the walk-in field cages, they generally landed on the supporting frame of the cage, its side walls and roof. The flies would clean themselves and occasionally fly short distances. The irradiated only (Irrd) males or the males that were irradiated and exposed to the release process at different durations (5m, 60m and 120m) competed successfully with the untreated colony males under our experimental field cage conditions. There were no significant differences in the relative mating index values between irradiated only males (Irrd) and the treatment males (**Table 2, Figure 5**). The relative mating indices showed that the untreated colony male flies do not outcompeted the male flies that were irradiated but not exposed to the automated chilled release process or the male flies that were loaded into the machine except 120m (G= 4.591. df=1, *P* = 0.032) (**Supplementary file 3**).

**Figure 5:**
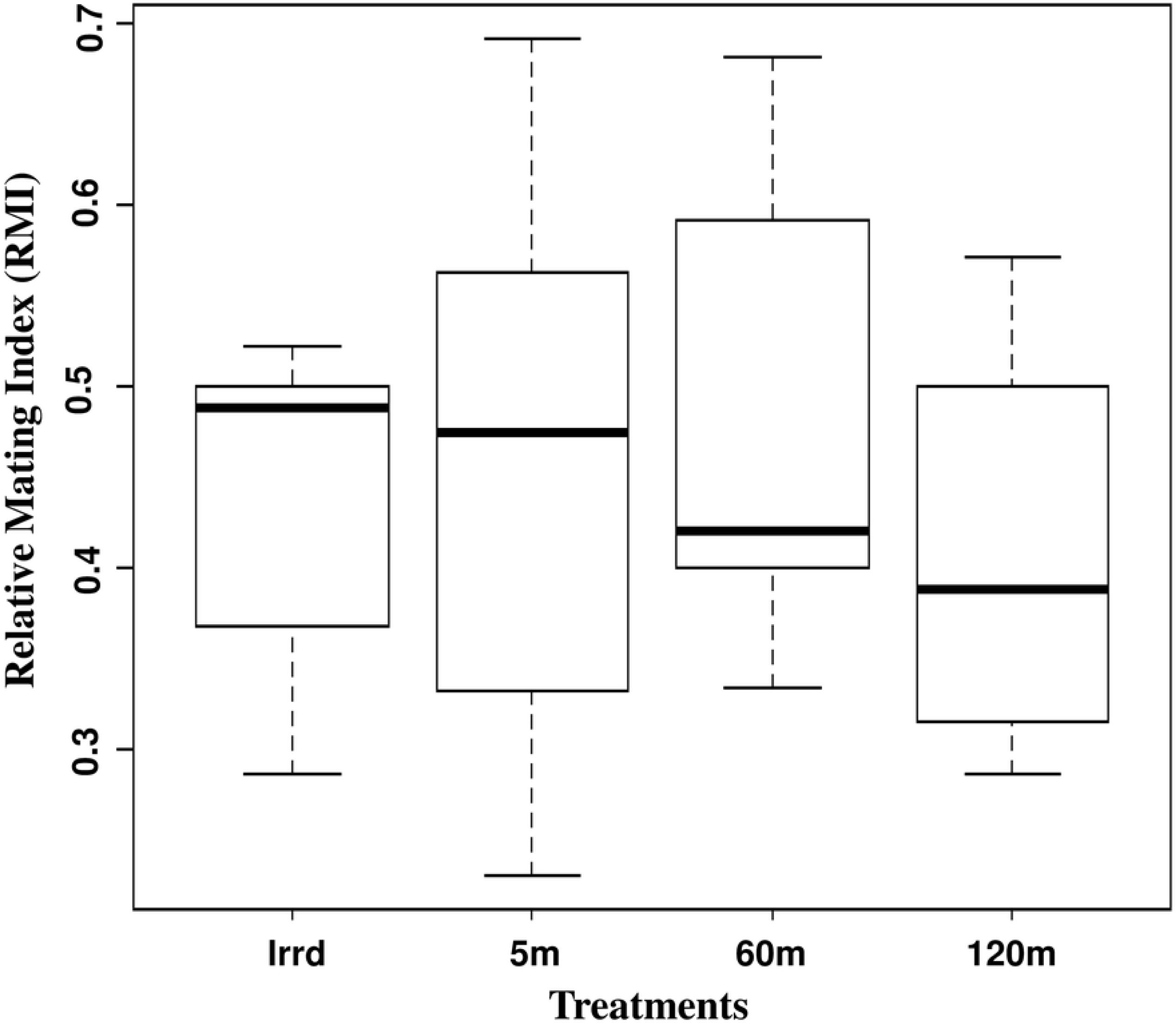
Impact of the release process through the BSI™ on male relative mating index (RMI). This graph represents the minimum, first quartile, median, third quartile and maximum for each treatment. Values indicated by the same lower-case letter do not differ significantly at the 5% level.

#### c) Pre-mating period and mating duration

Mating pairs were formed soon after the males were released in the field cages. There were no significant differences in the pre-mating period between the only irradiated males (Irrd) and the irradiated males exposed to the release process at different durations (5m, 60m and 120m). The premating period also did not differ between the different treatments (**Figure 6a**). Similarly, there were no significant differences in the mating duration between only irradiated males and the treatment males (5m, 60m and 120m) (**Figure 6b, Table 2**).

**Figure 6:**
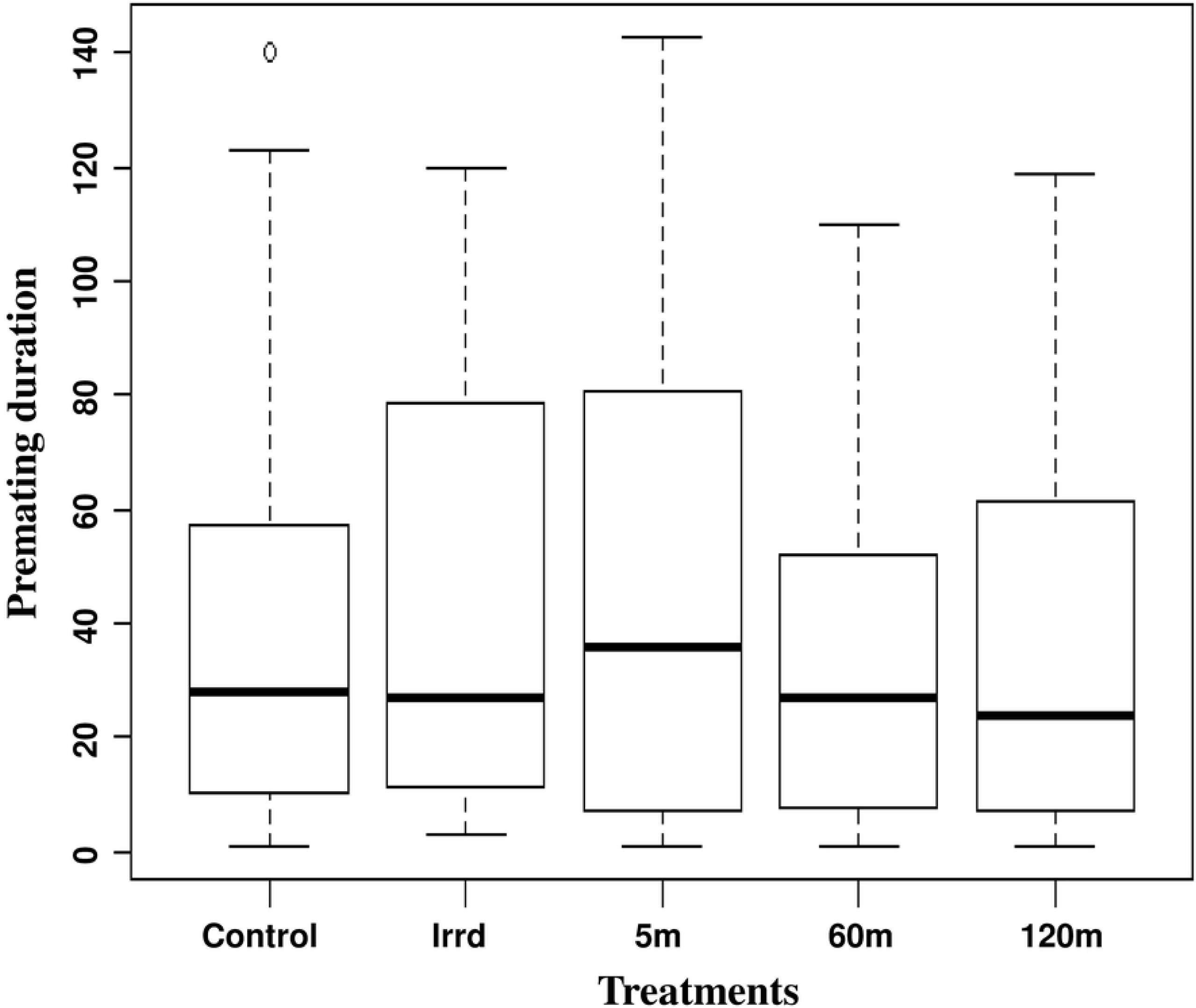
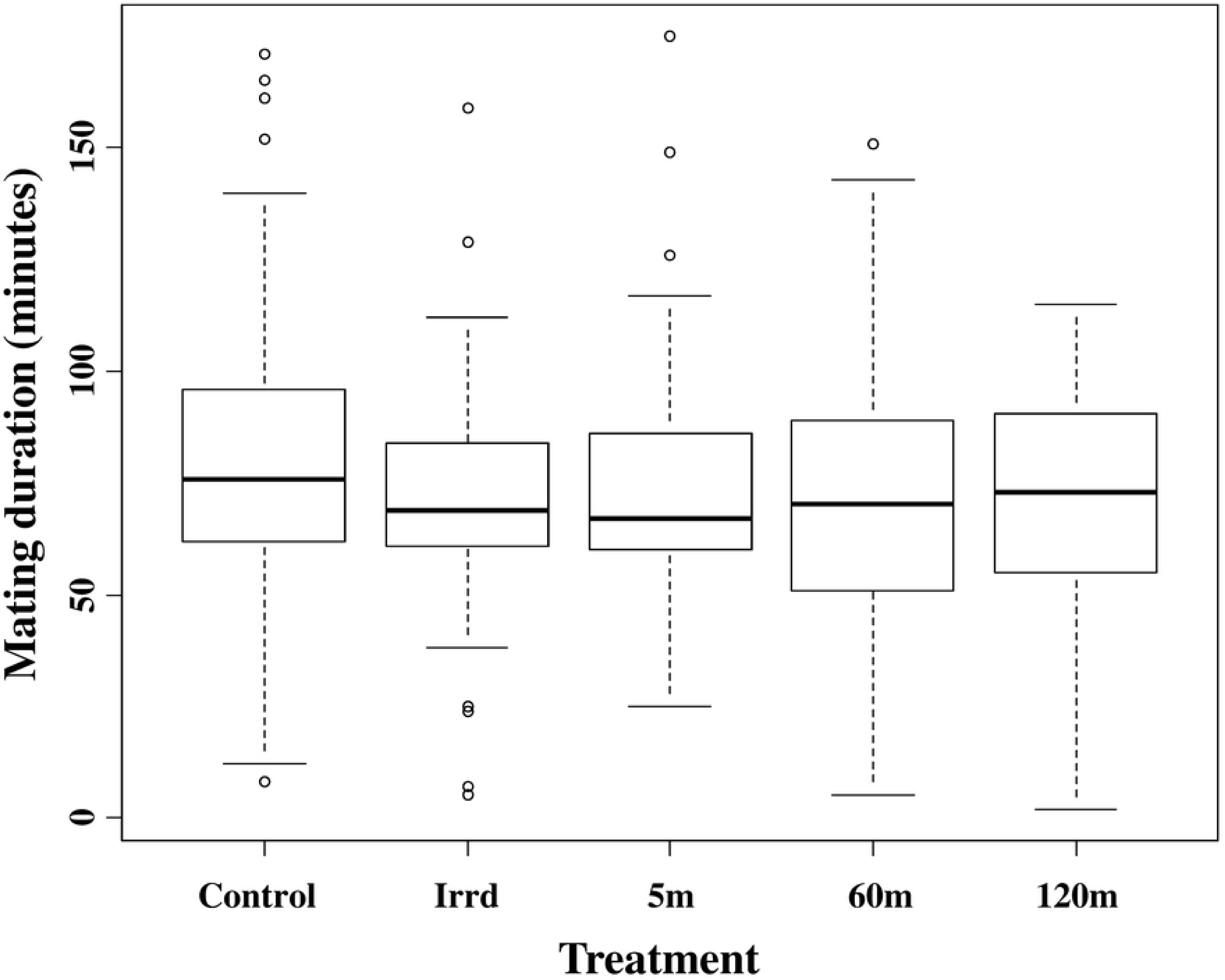
Impact of the release process through the BSI™ on male premating period (a) and mating duration (b). The graph represents the minimum, first quartile, median, third quartile and maximum for each treatment.

#### d) Insemination rate

Females mated with males of the different treatments had similar insemination rates (**Figure 7A**). In general, 55-69% of the dissected mated females showed spermatheca that were completely filled with sperm regardless of the treatment to which the males were exposed, while the rest of the dissected mated females had a similar distribution of empty and partially filled spermatheca (13-26%) **Figure 7B**).

**Figure 7.**
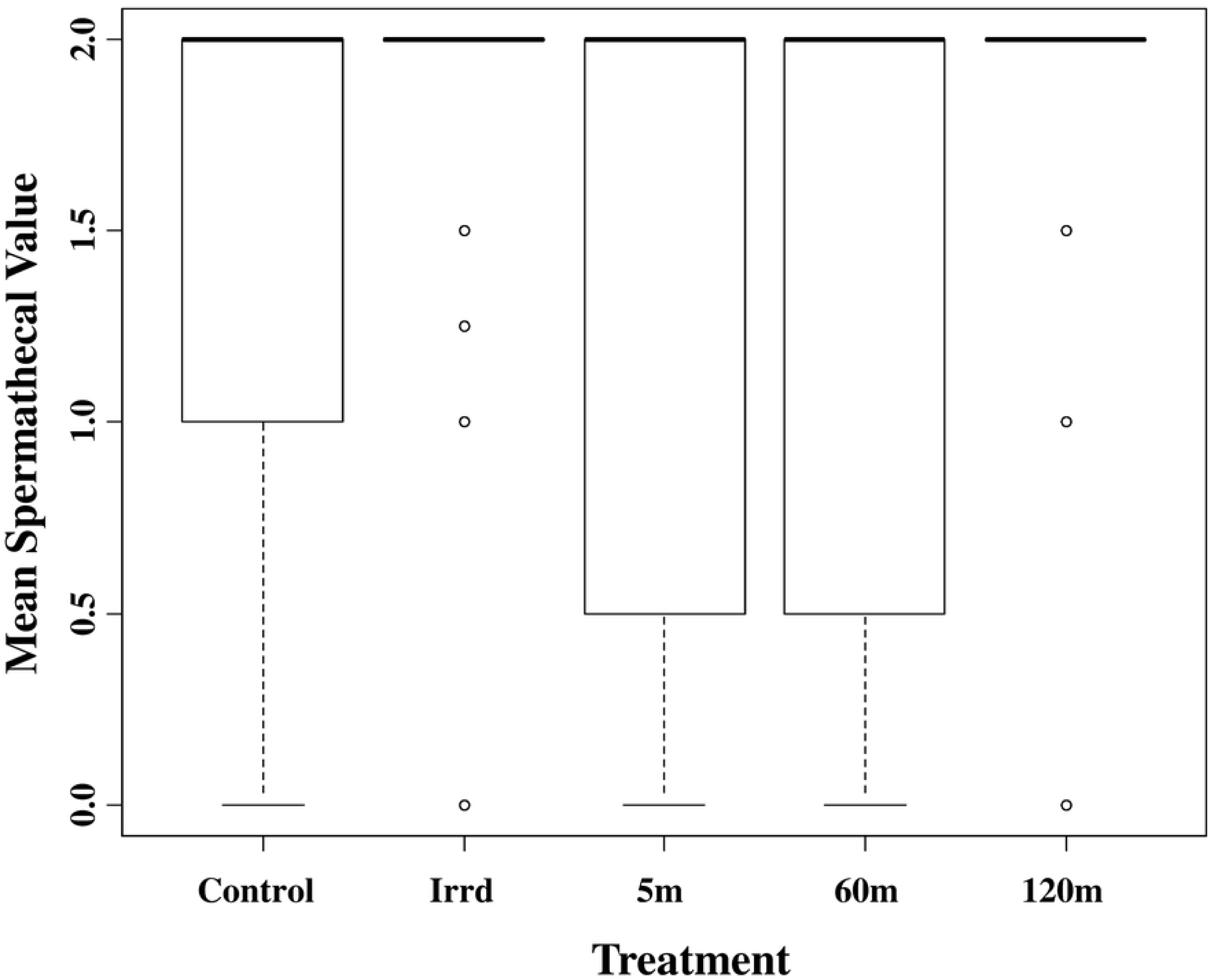
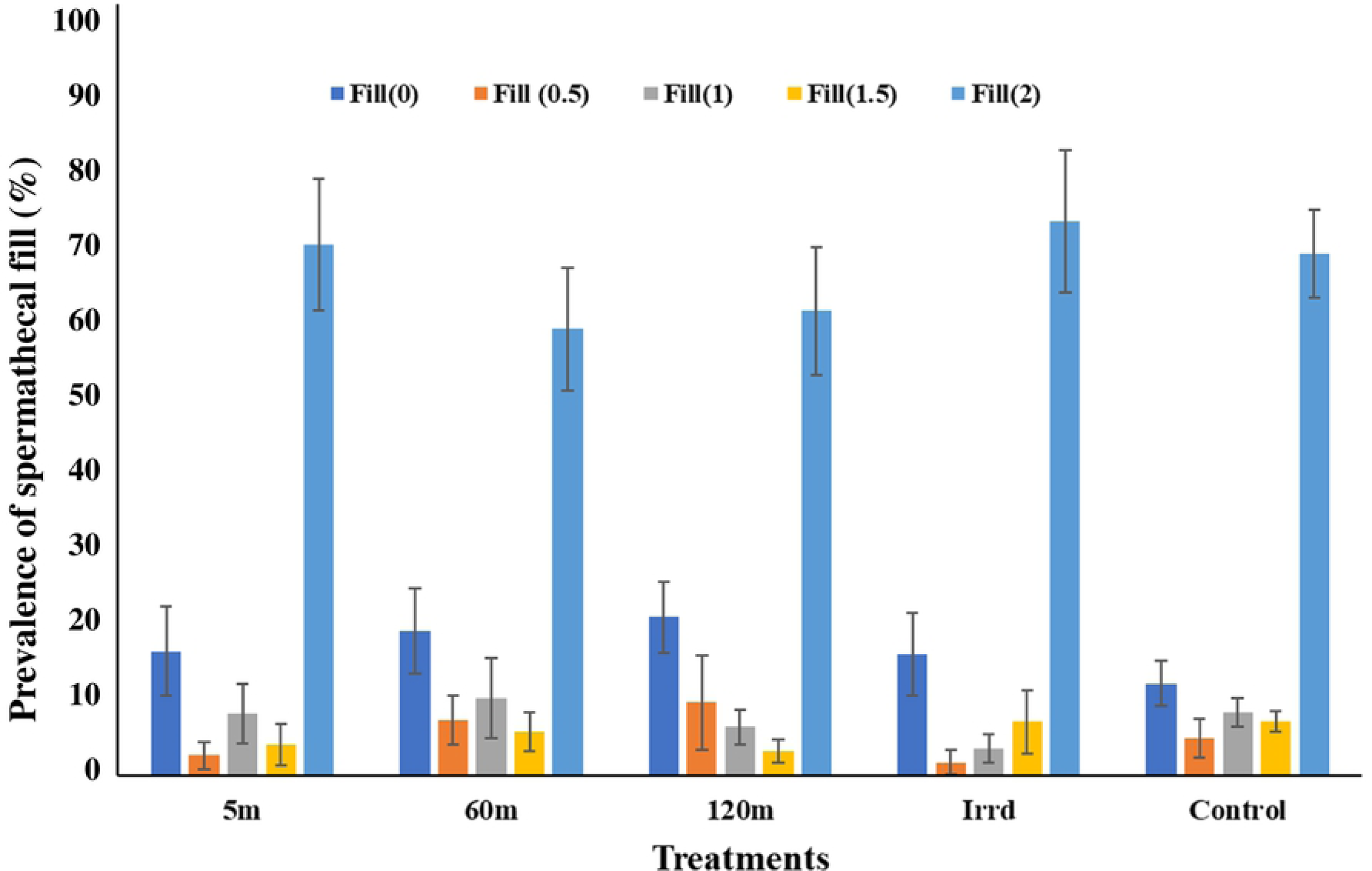
Impact of the release process through the BSI™ on the insemination rate of mated females a: Mean spermatheca value, b: distribution of spermatheca fill.

#### e) Male survival

The results from the Kaplan-Meier analysis indicate that under starvation conditions, most of the males died within two weeks but they survived relatively longer (50% of males survive > 20 days regardless treatments) when receiving a normal feeding regime of three blood meals per week (**Figure 8**). Survival of males chilled for 120 minutes before release and starved lived significantly shorter than only irradiated males (z = 2.954, *P* = 0.00313). However, no significant difference in male survival was observed between irradiated only males (Irrd) and those exposed to the release process for 5 or 60 minutes (**Figure 8A**). Under feeding conditions, males that were only irradiated (Irrd) died significantly faster than males of the 5 min treatment group (z = 2.90, P = 0.00369). However, no significant differences were observed between the irradiated only treatment (Irrd) and the 60m and 120m treatment groups (**Figure 8B**).

**Figure 8:**
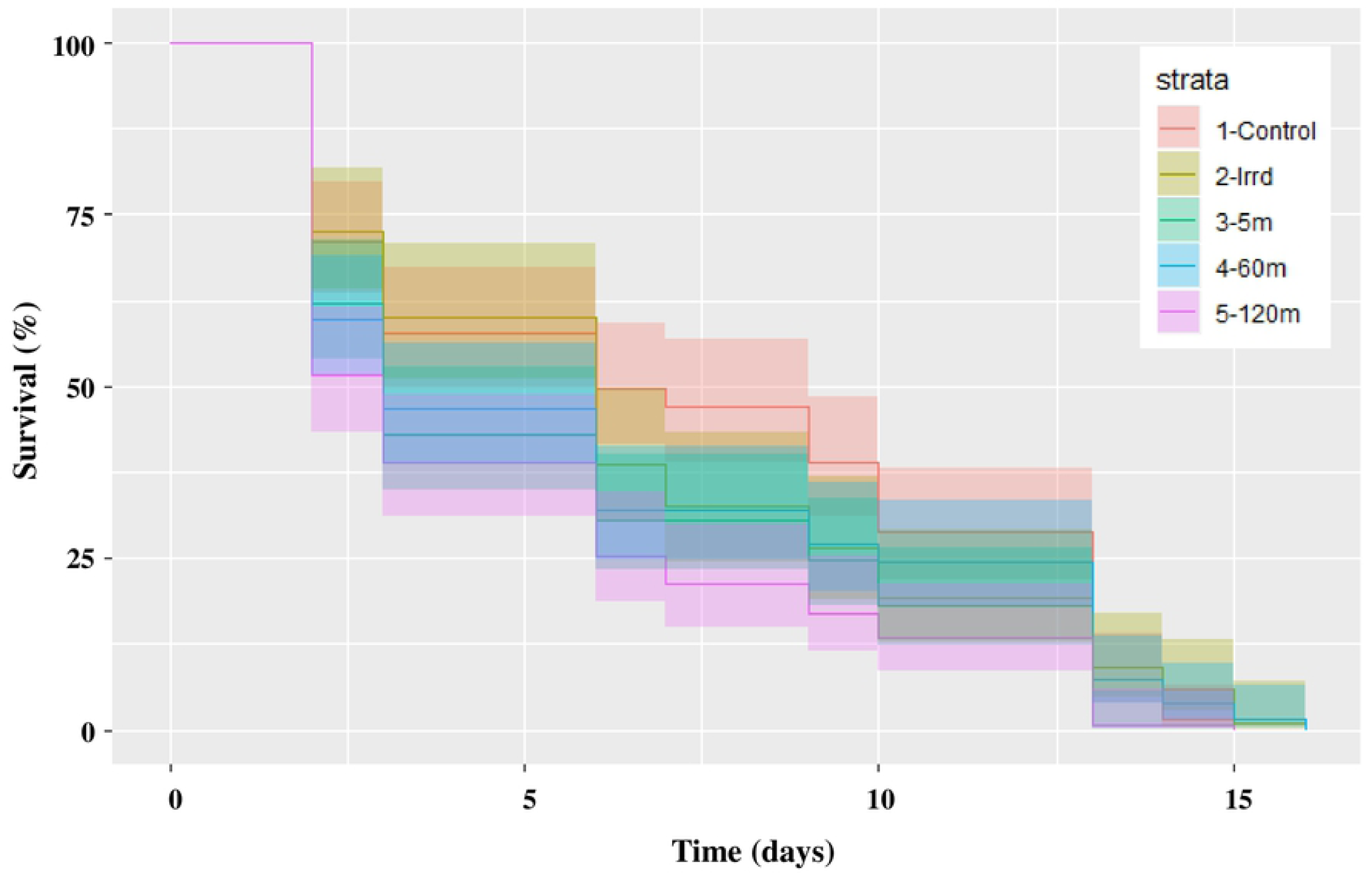
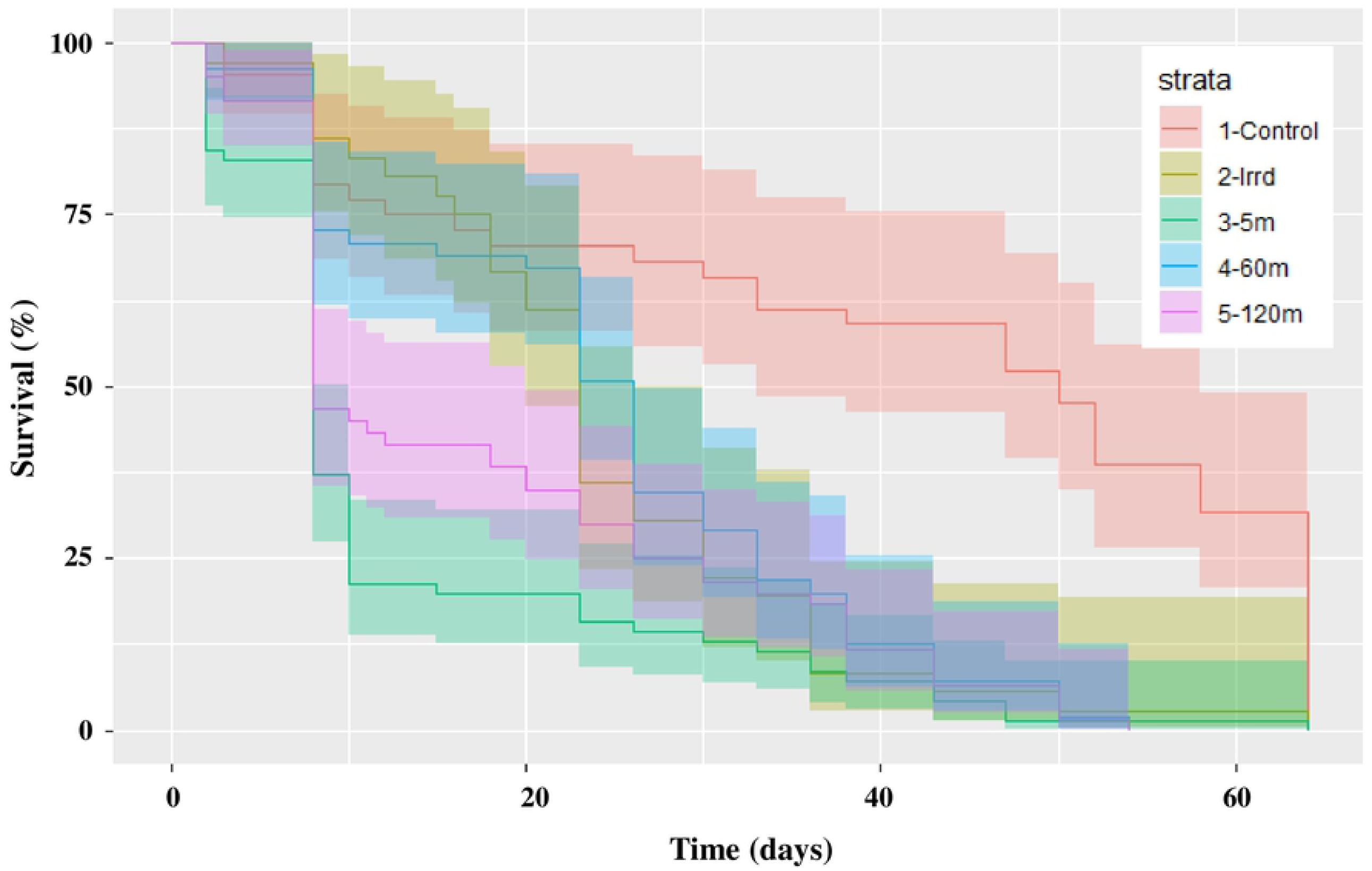
Impact of the release process through the BSI™ on *Glossina palpalis gambiensis* male survival under starvation (A) and feeding (B) conditions.

## Discussion

The main objective of this laboratory study was to evaluate a new prototype of an automated chilled adult release device (BSI™) to release sterile male tsetse flies from the air, firstly to assess its functionality and suitability for the release of sterile tsetse flies at low density (i.e. 10 males /km^2^) and secondly, to assess the impact of the release process under chilled conditions on the quality of the sterile males. Generally, during flight simulations in this study the BSI™ device released flies at a consistent release rate (60±9.58 males/min for the speed of 0.6 rpm), demonstrating its functional ability to homogenously release sterile tsetse flies in target areas (an important aspect of sterile insect release devices) [31]. The accuracy and consistency may be attributed to its release mechanism that consists of a rotating cylinder which rotates at a constant speed (0.6 rpm), allowing a loading of similar number of flies per unit time into the cylinder. This mechanism is different from the ones used in earlier release devices, i.e. a conveyor belt or a vibratory feeder used in the MSRM1 and MSRM2, respectively [37]. Although, it is noted that the MSRM2 fulfilled the requirement of the tsetse eradication program in Senegal for very low release rates (∼50 flies/km^2^) and was routinely used for some time to replace the release cartons, the machine was still not able to achieve the lowest rate of 10 flies/km^2^ without manipulation from the handlers and has been discontinued from use as previously mentioned [37].

The BSI™ is controlled by Bluetooth^®^ from a tablet computer that includes a completely automated guidance and navigation system, providing the pilot of the release aircraft with the polygon areas and the necessary data on the wild flies’ distribution and density on the ground. The control system is also able to display the release rates and the physical conditions in the machine such as temperature, relative humidity, speed and vibration power of the shaker. The automated navigation system gives an advantage in accuracy and homogeneity over earlier release systems such as the use of carton boxes that was prone to human error [37]. The observed accuracy and consistent release rates can also be attributed to the absence of clumping of flies in the machine during release regardless of the number of flies loaded into the device (max. 2000 flies). Clumping was prevented by maintaining the flies in a free-flowing granular state at a temperature of 6 ±1°C, an improvement as compared with the previously used MSRM2 machine where clumping was a serious problem [37]. The BSI™ has several advantages compared with the MSRM2 machine in terms of weight (20 kg versus 64 kg), power (2-3A −12V versus 100 A), and it is less bulky. However, like the MSRM2 device, it can be easily installed in a gyrocopter **(Figure 1b)**. These advantages of the BSI™ device in terms of its efficient release mechanism, its automated navigation system, its low weight and power requirements and the absence of clumping of the flies, will make it an attractive option for operational use in field release programs. It can be operated solely by the pilot, and hence eliminates the necessity of a release coordinator, reducing the costs of tsetse fly aerial release [37].

Generally, the results show that the combined effect of chilling and the mechanical abrasion experienced by the flies when passing through the release device did not have a significant negative impact on flight ability, relative mating index, premating and mating duration of the flies as compared to the irradiated flies. In this case, the absence of a negative effect may have been contributed by shorter chilling conditions that did not exceed three hours (including the handling procedure before the longest release period). In addition, the insemination rates in female flies that had mated with males that had been chilled in the machine for at least an hour before being released, were not significantly reduced. This does not corroborate the results of the Mutika et al [43] study that simulated long distance transport of pupae and the release of *G. p. gambiensis* as chilled adults (pupae stored for 5 days at 10°C and sterile males stored up to 30 h at 5.1 ± 0.4°C), i.e. prolonged chilling of adults affected the biological quality of the flies, and the study recommended that the duration of chilling should be minimized. The significant reduction in flight ability regardless of the chilling and release process) of irradiated males as compared with the control males (not irradiated, not passed through the machine indicates a significant negative impact of irradiation on the released males. These results are in agreement with many previous studies that report a dose dependent negative impact of irradiation on male insect quality [58–60]. In addition, these results are in agreement with those of Diallo et al. [61], who found that irradiation combined with chilling conditions increased flight propensity in comparison with irradiation alone. Overall, our study further agrees with that of Mutika et al [44] on the combined effect of irradiation and chilling in that a significantly lower proportion of sterile males stored at low temperatures succeeded in securing mates compared with untreated males. Generally, irradiation only did not affect the survival of starved male flies but the release process at 120 min significantly reduced their survival. The lower survival observed in our study is in agreement with Mutika et al. which clearly stated that the longer the adults are kept at the low temperatures, the lower their biological quality [43]. Also, this result agrees with the study of Diallo et al. [61] in which starved sterile males that emerged from chilled pupae showed an average survival of 4-5 days. Similarly, under feeding conditions, the males released after a short duration within the machine (5 min) died significantly faster than the irradiated (Irrd) and non-released males (control). Surprisingly, males released after a longer time in the machine survived better, which might be due to a reduced metabolic rate due to the chilling that could have stimulated cellular repair mechanisms of the somatic damage [61].

Despite the encouraging results, the BSI™ has limitations that ideally need to be improved. First, there were errors in fly counts that were related to more than one fly at a time being loaded into the cavity of the rotating cylinder. This can somehow be corrected as the BSI™ software allows a factor to be included to correct errors in counts (as displayed on the navigation page during the flight simulations). This error might be related to the cavity size and tests should be conducted to assess the effect of different (reduced) sizes of the cavity to receive only one fly at a time. However, even after the correction of errors the software estimated lower maximum counts (ca. 1598 (**Figure 3A**)), compared with the 2000 flies loaded into the machine. This estimation may be improved by implementing the statistical model used in this study (**Figure 3B**) to give more accurate predictions of the real counts. Additionally, handling (counting, sorting) of flies before the release process could have contributed to the overall performance of the flies because of the increased length of time under low temperatures. In future, with new technologies that are currently under research [62,63], male and female pupae can be separated on day 23-24 post larviposition, hence eliminating the counting and sex sorting steps before loading into the BSI™ as done in this study as well as the transport under chilling which was not part of this study. In addition, the counting could also be eliminated by using weight or volume to estimate the number of flies, especially where large numbers of flies will need to be loaded into the machine during SIT programs.

The sterile males released in Senegal are produced in tsetse mass-rearing facilities located in other countries (i.e, The Slovak Academy of Sciences, Slovakia, the IPCL, Seibersdorf, Austria and the Insectary of Bobo Dioulasso (IBD) and CIRDES, Bobo Dioulasso, Burkina Faso) and require transport under chilled conditions (10°C) to prevent emergence during shipment [29,41,61,64]. Although in this study the male pupae were chilled at 10°C for 1-2 hours before irradiation, this treatment did not accurately mimic what occurs in the operational release program in Senegal, i.e. the pupae in the study were not packaged or shipped and the chilling time of the adult flies at 10°C was relatively short. Further studies are needed to assess the combined effect of chilling, packaging and shipment with the release process using the BSI™. In addition, the distribution and recapture of released males under field condition remains to be assessed.

In conclusion, in an effort to expand the repertoire of machines available for tsetse release in the field and improve previously tested release mechanisms, our results show that the BSI™ has great potential that merits consideration for use in the current SIT release program in Senegal.

## Acknowledgments

We thank technical staff of the Insect Pest Control Laboratory, Seibersdorf for tsetse fly maintenance. We thank Drs Hamidou Maiga and Güler Demirbas-Uzel for their help in the statistical analysis. This work was funded by the Joint Food and Agricultural Organization of the United Nations (FAO) / International Atomic Energy Agency (IAEA) Division of Nuclear Techniques in Food and Agriculture and by the European Research Council under the European Union’s Horizon 2020 research and innovation programme (grant agreement No 682387—REVOLINC). This paper reflects only the authors’ views and the IAEA is not responsible for any use that may be made of the information it contains.

## Author Contributions

Conceived and designed the experiments: AMMA AGP MJBV JB JBr. Performed the experiments: CKM GNM. Analyzed the data: AMMA AGP JB GNM. Contributed reagents/materials/analysis tools: CKM AMMA AGP MJBV JB. Wrote the paper: CKM GNM AMMA AGP MJBV JB EB JB MTS BS MVO.

## Conflicts of interest statement

Jimmy Bruno designed the release system (Bruno Spreader Innovation, (BSI™) and works for his own enterprise Jimmy Bruno, AEWO (aerial works).

## Availability of data and materials

Materials described in the manuscript, including all relevant raw data, are available in this link https://dataverse.harvard.edu/dataset.xhtml?persistentId=doi:10.7910/DVN/AQIW8P.

